# Odor-evoked respiratory responses throughout development in sighted and blind mice

**DOI:** 10.1101/2025.02.18.638853

**Authors:** N. Bouguiyoud, P. Litaudon, J. Frasnelli, S. Garcia, B. Messaoudi, AM. Mouly, S. Al Aïn, E. Courtiol

## Abstract

Congenital blindness affects olfactory function depending on developmental stage. However, when studying the ontogeny of olfactory abilities, not all behaviors are expressed at every age making the longitudinal comparisons difficult. Odor-evoked respiratory responses, which are unlearned and do not require complex motor coordination, may serve as sensitive measures of olfactory abilities throughout ontogeny. Using a non-invasive measure of respiration in an olfactory perceptual paradigm, we assessed odor-evoked respiratory responses in a model of congenital blindness at 3 ages, infant, juvenile and adult, in the same mice and in both males and females. We demonstrated the differential outcome of two respiratory parameters (i.e. frequency and amplitude) in a mouse model of congenital blindness. We showed that blind mice have similar olfactory abilities than sighted mice throughout ontogeny but display enhanced sniffing frequency and amplitude, starting at the juvenile age for the latter one, that may help them better explore their environment. We also demonstrated that respiratory frequency is a robust index of age and of olfactory detection, habituation and discrimination at all ages. On the other side, respiratory amplitude does not provide a proxy of olfactory performance at all ages, however, it does highlight differences between sexes and phenotypes associated with visual deprivation. To conclude, our data highlight that respiratory parameters can be used as a complementary approach to assess olfactory performance throughout development and provide an index of olfactory plasticity.

## Introduction

Early visual deprivation enhances the remaining non-visual sensory modalities (Kupers and Ptito 2014) such as audition and touch (Van Boven et al. 2000; Voss et al. 2004). Similarly, compared to sighted mice, blind rodents exhibit superior olfactory function when measured behaviorally (Bouguiyoud et al. 2022; Touj et al. 2020; Zhou et al. 2017). Congenital blindness affects olfactory function depending on developmental stage, as odor-guided behaviors do not differ in infants (PN7), but become apparent in juvenile (PN33-36) and adult mice (Bouguiyoud et al. 2022). However, the same behavioral paradigms cannot be applied in all age groups. Nevertheless, one potential method to assess olfactory function across the lifespan is to measure respiration.

Respiration is a highly dynamic motor act (Freeman et al. 1983; v. Holst and Kolb 1976; Moulton and Marshall 1976; Rehn 1978; Teghtsoonian and Teghtsoonian 1982; Teichner 1966; Walker et al. 1997; Welker 1964; Wesson et al. 2008b, 2008a, 2009; Youngentob et al. 1987) and is typically measured by assessing respiratory frequency (Youngentob et al. 1987). Respiration is influenced by vigilance (Bagur et al. 2021; Girin et al. 2021; Janke et al. 2022; Zeng et al. 2012), emotional state (Dupin et al. 2019, 2020; Hegoburu et al. 2011; Moberly et al. 2018), metabolic status (Aimé et al. 2012; Al Koborssy et al. 2019; Tong et al. 2011), and motivation (Clarke and Trowill 1971). In other words, respiration measures inform on the internal state. Consequently, respiratory parameters can also accurately be used to distinguish behaviors (e.g. immobility, freezing, sleep) through supervised classification (Janke et al. 2022).

One particular form of respiration is sniffing, i.e., a regular succession of high frequency respiratory cycles combined with stereotyped movements of the vibrissae, nose and head (Deschênes et al. 2012; Welker 1964; Youngentob et al. 1987). Respiration/sniffing and olfaction are inextricably linked as respiration is the vector of odorant molecules. In fact, sniffing allows for active odor sampling (Wachowiak 2011; Welker 1964). Respiration patterns depend on and are strongly modulated by olfactory tasks (Courtiol et al. 2014; Hegoburu et al. 2011; v. Holst and Kolb 1976; Kepecs et al. 2006, 2007; Lefèvre et al. 2016; Reisert et al. 2020; Rojas-Líbano and Kay 2012; Wesson et al. 2008a, 2008b). Further, odor-evoked sniffing is a more robust and sensitive measure of olfactory abilities than classical measures of exploration (Coronas-Samano et al. 2016; Johnson et al. 2020a; Wesson et al., 2008).

Remarkably, odor-evoked respiratory responses, which are spontaneous, unlearned and do not require complex motor coordination (Boulanger-Bertolus et al. 2023), may allow to compare olfactory abilities throughout development. In rats, sniffing reflects olfactory abilities early in rat development (Alberts and May 1980a, 1980b; Boulanger Bertolus et al. 2014; Boulanger-Bertolus et al. 2023; Seelke and Blumberg 2004). Indeed, already newborns (at post-natal day 1: PN1) (Alberts and May 1980a, 1980b; Boulanger-Bertolus et al. 2023) and sleeping infants (PN8) (Seelke and Blumberg 2004) respond to odorants with an increase in respiratory frequency. Although in rats stereotyped sniffing behavior only starts at PN11, sniffing bouts can already be observed around PN3 (Alberts and May 1980a), while high-frequency sniffing (>4Hz) is established at PN15 (Zhang et al. 2021). As early as PN7-10, respiration modulates activity in olfactory processing centers (Fletcher et al. 2005; Gretenkord et al. 2019), although high frequency oscillations emerge only around PN15-16 along with the more robust expression of high-frequency sniffing (Zhang et al. 2021). Despite this plethora of studies, only one study directly compared odor-evoked respiratory responses between different developmental stages, from infancy to adulthood (Boulanger-Bertolus et al. 2023). In contrast to rats, the development of odor-evoked respiratory responses in mice are still relatively unknown. Therefore, we not only aimed to assess odor-evoked respiratory responses across the lifespan in sighted mice but also to use those responses to assess olfactory functions in congenital blind mice across different developmental stages. To do so, we used an odor habituation/cross-habituation task at three ages on the same animal: infant (PN10-13); juvenile (PN30-34) and adult (PN>60). Odor habituation/cross-habituation paradigm is a useful paradigm, applicable at all ages, that allows the study of odor detection, habituation, discrimination and short-term memory processes, and in which sniffing frequency has been shown to be a useful and sensitive readout of olfactory abilities (Al Koborssy et al. 2019; Coronas-Samano et al. 2016; Johnson et al. 2020; Walker et al. 2024; Wesson et al. 2008b). In addition, given the importance of other respiratory parameters (Youngentob 2005; Youngentob et al. 1987), the respiratory cycle total amplitude was also analyzed. We used a murine model of congenital blindness, the ZRDBA mouse strain (Touj et al. 2019; Tucker et al. 2001). It generates, within the same litter, sighted and blind pups, with the latter being characterized by an absence of eyes, optic tracts, and retina-hypothalamus afferents.

## Material and Methods

### Animals

The experiments were carried out according to the ethical guidelines of the European Communities Council Directive 2010/63/EU, as well as the approval # 34624 of the CELYNE ethical committee and the Ministry of Higher Education, Research and Innovation. All efforts were made to minimize the number of animals used by performing a longitudinal study on the same animal. Each experiment was designed in agreement with the “3R” rules.

Thirty 6-9 weeks old ZRDBA mice (10 blind females, 10 sighted females, 5 blind males and 5 sighted males) generated at the University of Québec in Trois-Rivières, were transported to the Lyon Neuroscience Research Center Animal facility. These breeders were used to launch a line of ZRDBA mice in Lyon. In this strain, half of the littermates homozygous for the Rx/Rax gene (chromosome 18; (Tucker et al. 2001)) are born blind and the other heterozygous half are born sighted (Bouguiyoud et al. 2022a;b, 2023; Touj et al. 2020).

A total of 11 litters were used with a total of 46 animals including 11 blind males, 12 blind females, 12 sighted males and 11 sighted females. We had technical issues with one blind male and one sighted male during on one of the behavioral sessions; we therefore did not consider them for the analysis leaving a sample of 10 blind males, 12 blind females, 11 sighted males and 11 sighted females (Fig. 1A). Animals were housed under controlled optimum conditions including temperature 22 ± 2 °C, hygrometry 50 ± 10%, constant light / dark cycle 7:00 am to 7:00 pm, with food and water *ad libitum*.

**Figure 1.**
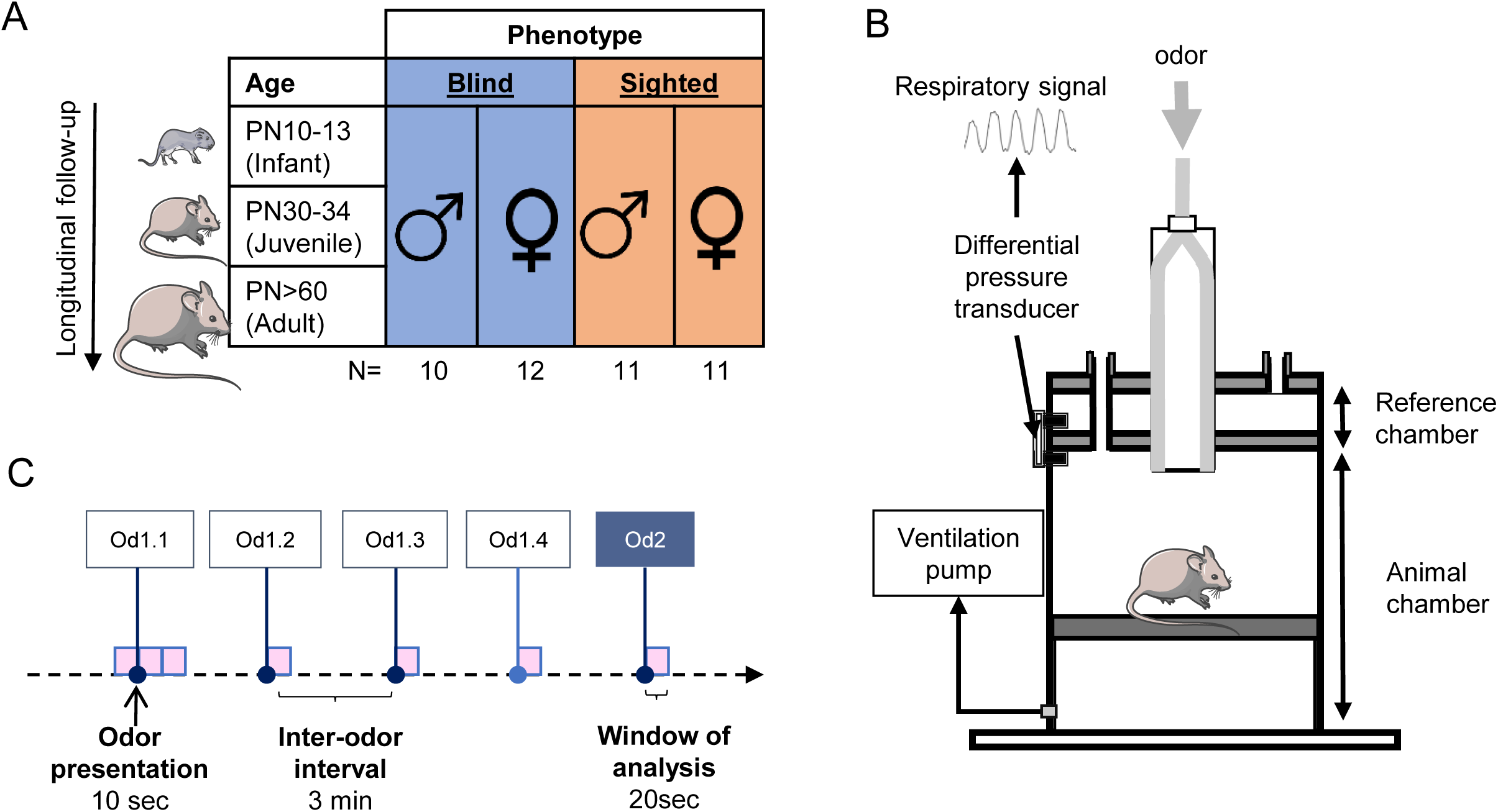
A- Each mouse was tested at three different ages : infant (PN10-13); juvenile (PN30-34) and adult (PN>60). We had four groups : blind males (n=10); blind females (n=12); sighted males (n=11); and sighted females (n=11). B- Design of the whole-body plethysmograph, with the animal chamber and the reference chamber. Clean air and odors arrived through the top of the cage and air was vacuumed out at the bottom. C- Odor habituation/ cross habituation protocol : Od1 is presented four times followed by the presentation of Od2. Each odor is presented during 10 sec and there is an inter-odor interval of 3 min. The time-window of respiratory signal analysis was 20 sec (pink squares). Each animal underwent this paradigm at each age with the following odor pairs : 1.8 cineole/ ethyl-heptanoate at the infant age, linalool / anethole at the juvenile age and peppermint / limonene at the adult age.

### Respiration recordings

To comply with the reduction principle of the 3R, longitudinal measurements in each mouse were performed at three different ages: infant (post-natal day 10-13; PN10-13) when their eyes were either still closed or opened very recently; juvenile (PN30-34) and adult (PN>60; Fig. 1A).

#### Apparatus

The apparatus consisted of a whole-body plethysmograph [diameter : 20 cm and height : 16.5 cm; EMKA Technologies, France; Fig. 1B; (Boulanger-Bertolus et al. 2014; Courtiol et al. 2014; Lefèvre et al. 2016; Boulanger-Bertolus et al., 2023)]. The whole-body plethysmograph allows non-invasive recording of respiration of unrestrained animals. It consists of a plexiglass cage composed of two independent compartments: the recording one where the mouse is installed and the reference one (Fig. 1B). The changes in pressure resulting from the mice’ respiration were detected using a differential pressure transducer located between the recording compartment and the reference one. The signal was sampled at 1 kHz, amplified (Amplipower, EMKA Technologies) and recorded using a custom-made software developed in Python (pyacq, https://github.com/pyacq/pyacq). The height of the plethysmograph was adapted for the infants in order to optimize the signal to noise ratio by adding a custom-made design piece of polystyrene to reduce total volume. The infant age (PN10-13) was also chosen to have a sufficient signal to noise ratio with our apparatus. At this age, a heating pad was also used beneath the whole-body plethysmograph to increase the temperature for the well-being of PN10-13 pups.

Calibration of the device was performed three consecutive times by injecting 1 ml of air through a syringe at the bottom of the whole-body plethysmograph. The average of all 3 values was calculated using our custom-made software to apply a calibration factor prior to each mouse and each session.

The whole-body plethysmograph was placed in an artificially illuminated sound-attenuating enclosure, isolating the animal from the surrounding environment (L 60 cm, W 60 cm, H 70 cm). The entire experiment was video recorded via two webcams (HD Webcam Logitech) mounted on the side wall of the sound-attenuating enclosure. Videos were captured using the pyacq software and synchronized to the respiratory signal for each session.

A constant airflow was delivered from the top of the whole-body plethysmograph (2l/min; Fig. 1B). The air was vacuumed out at the bottom at 2l/min to ensure constant ventilation. During the olfactory task, pre-selected pairs of odors were presented via a custom-made olfactometer through the main air stream. The odor was controlled with a solenoid valve that diverted the airflow to the odor tube, thus minimizing pressure change (2l/min; 1:8 vapor to air).

#### Odor habituation/cross-habituation

At each age (infant, juvenile and adult), mice were placed in the experimental chamber for 20 minutes and odors were presented in a habituation / cross-habituation paradigm (Al Koborssy et al. 2019; Walker et al. 2024).

After three minutes in the plethysmograph, a first odor (Od1) was presented four times (Od1.1, Od1.2, Od1.3 and Od1.4), followed by the presentation of a second odor (Od2). Each odor was presented during 10 seconds thanks to an automatic setup, with an inter-odor interval of 3 minutes (Fig. 1C). For our study, all odorants used were unfamiliar to the animals. The pairs of Od1/Od2 odors were different at each age in order to avoid any bias related to residual learning, with 1.8 cineole / ethyl-heptanoate at the infant age, linalool / anethole at the juvenile age and peppermint / limonene at the adult age (odors from Sigma-Aldrich).

### Data analysis

#### Respiratory signal

The analysis of the respiratory data relied on a Python-coded algorithm allowing the detection of the transition point between inspiration and expiration (Courtiol et al. 2014; Girin et al. 2021; Lefèvre et al. 2016; Roux et al. 2006; Walker et al. 2024). The signal was first processed with a high-pass filter at 2 Hz. Inspiration was defined to start at the zero-crossing point of the falling phase and ends at the zero-crossing point of the rising phase, whereas the expiratory phase began at the zero-crossing point of the rising phase and ends at the zero-crossing point of the falling phase. Following the detection of each respiratory cycle, values of respiratory cycle parameters were extracted. We notably analyzed the instantaneous frequency (1/cycle duration; Hz). In addition, we analyzed the total amplitude, representing the sum of the expiratory and inspiratory peak flow rates, normalized to the weight of the animal (ml/sec/g) as this measure strongly depends on the weight of the animal (Mortola et al. 1994).

In order to reduce noise and to eliminate potential artifacts, raw data were visually inspected, and an automatic Python script was further used to remove artifacts from the dataset. Cut-off values were determined upon the 5 first minutes of each recording. The median and median absolute deviation of the total amplitude were estimated for each mouse and session over that 5-minute period. Each cycle for which value was above the threshold of median + 8x median absolute deviation was discarded. When artifacts were separated by less than a 2-sec period, neighboring cycles were agglomerated into batches which were also discarded.

Using the odor habituation/cross-habituation paradigm, we assessed 1) the odor-evoked sniffing in response to the first presentation of an odor, 2) how this response evolves over repetition of that same odor and 3) the sniffing response to a new odor (Al Koborssy et al. 2019; Walker et al. 2024). This allowed us to test for detection, habituation, discrimination of an odor and short-term memory processes.

The instantaneous respiratory frequency was first averaged on a sliding window, with local gaussian kernel of 1-sec bin, leading to 1-sec time bin curves before and after the odor onset (Figure 2A-B). The resulting individual curves were then averaged among animals of the same experimental group 40 sec before and 40 sec after the odor onset (Figure 2B).

**Figure 2.**
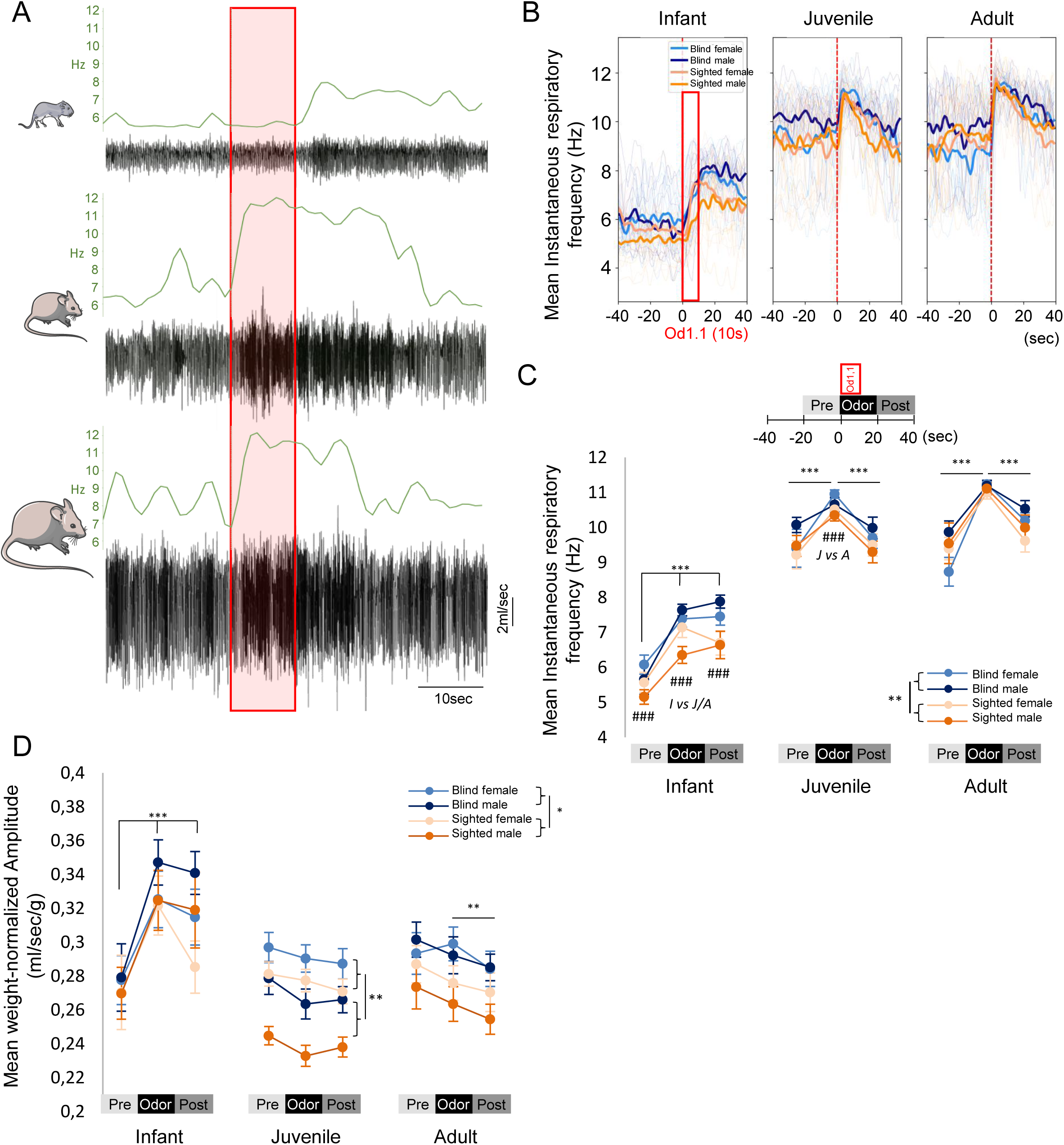
Respiratory responses to the first presentation of an odor. A- Example of sniffing response to a new odor in the same animal at each developmental age (from top to bottom: infant, juvenile and adult). Respiratory signals are in black; the instantaneous respiratory frequency is averaged on a sliding window, with local gaussian kernel of 1-sec bin, in green. The red outline represents the odor. B- The instantaneous respiratory frequency was first averaged on a sliding window, with local gaussian kernel of 1-sec bin. The resulting individual curves (fine lines) were then averaged among animals of the same experimental group (blind female: cyan n=12; blind male: dark blue n=10; sighted female: salmon n=11 and sighted male: orange n=11) 40 sec before and 40 sec after the odor onset for each developmental age (in bold). The red rectangle represents odor 1 presentation. C-D Mean values of instantaneous respiratory frequency (Hz; C) and mean weight-normalized amplitude of the respiratory signal (D) were averaged over 20sec-time window prior to odor onset, overlapping odor presentation and 20sec post odor-bin. Values were averaged per group and per age; Mean +/- SEM. Repeated measures ANOVAs were performed. The ANOVAs were followed by Tukey post-hoc comparisons when significant.*, p<0.05; **, p<0.01; ***, p<0.001. * represents differences between odor periods or sex at each age. It also highlights difference between blind and sighted mice no matter the odor periods, age and sex. For respiratory frequency, # represents a significant difference between ages at each period (no matter the group), ###, indicates p<0.001.

Respiratory signals were then analyzed and averaged over a 20-sec window : 1) prior to the first presentation of odor 1 (Pre-Odor; Fig. 1C), 2) following each odor presentation (Odor, the odor lasts 10 sec but does not vanish at 10 sec exactly), and 3) after this 20sec-odor presentation bin (Post-Odor). Mean values of instantaneous respiratory frequency (Hz) and total amplitude were calculated per animal and per period over these 20sec-time windows.

### Statistical analysis

All statistical analyses were performed using Jamovi (https://www.jamovi.org).

For the detection of the first odorant presentation (Od1.1), respiratory frequency and weight-normalized amplitude were analyzed during the Pre-odor period, the Odor period and the Post-odor period (Fig. 1C). Repeated measures ANOVAs were used. We had two between-subject factors : *phenotypes* (blind vs sighted) and *sexes* (male vs female) and two within-subject factors : *age* (Infant, juvenile and adults) and Od1.1 *periods* : (pre-Od1.1; Od1.1; post-Od1.1). The ANOVA was followed by Tukey post-hoc comparisons when significant.

In the odor habituation/cross-habituation paradigm, respiratory frequency and weight-normalized amplitude were analyzed following each odor presentation (Od1.1, Od1.2; Od1.3; Od1.4 and Od2; Fig 1.C). Repeated measures ANOVAs were used with the following variables: *phenotypes* (blind vs sighted) and *sexes* (male vs female) as between-subject factors, and *age* (Infant, juvenile and adults) and *odor presentations* (Od1.1; Od1.2; Od1.3; Od1.4 and Od2) as within-subject factors. The ANOVA was followed by Tukey post-hoc comparisons when significant. Results are presented as mean ± SEM. P values smaller than 0.05 were considered significant.

## Results

### Temporal course of sniffing in response to the first presentation of an odor throughout development in both blind and sighted mice

We first assessed the temporal course of the odor-evoked respiratory responses to the first odor presentation (Od1.1) at each developmental age (Figure 2) using both respiratory frequency and amplitude. Raw respiratory signals from the same animal at the infant, juvenile and adult age along with the individual binarized measures of the instantaneous respiratory frequency are presented in Figure 2A-B.

The instantaneous respiratory frequency was averaged over a 20sec time-window before (pre), during (Odor) and after Od1.1 presentation (Post; Fig. 2C). We observed a significant effect of *phenotype* (F(1,40)=10,38, p=0.0025). Blind mice displayed higher respiratory frequency than sighted mice. Further, we observed a significant effect of *age* (F(2,80)=396,24, p<0.001), an effect of the *odor periods* (F(2,80)=88,49, p<0.001) and an interaction between *age* and *odor periods* (4,160)=10,49, p<0.001). To disentangle this interaction, we carried out Tukey post-hoc tests.

Regarding the dynamic of the response per *age* group, we observed a significant increase between the pre-odor and the odor periods at all ages (for the 3 ages: pre vs odor, p<0.001). At the infant age, the respiratory frequency during the post-odor period remained similar to the odor period (pre vs. post: p<0.001; odor vs post: p=0.99). At the juvenile and adult ages, however, the respiratory frequency decreased significantly during the post-odor period compared to the odor period and returned to the pre-odor level (juveniles: pre vs post: p=0.99; odor vs post: p<0,001; adults: pre vs post: p=0.11; odor vs post: p<0.001).

Regarding the interaction of *age* * *odor periods*, the mean pre-odor respiratory frequency was smaller in infants (5.6 Hz) than in both juveniles and adults (respectively 9.5 Hz and 9.4 Hz), and not different between the two latter (Infants vs juveniles/adults: p<0.001; adults vs juveniles: p=0.99). During the odor period, odor-evoked respiratory frequency significantly increases with ages (from 7.1Hz to 10.6 to 11.1 for infants, juveniles and adults, respectively; for all comparisons, p<0.001). During the post-odor period, respiratory frequency also significantly increases between infants and both juveniles and adults (p<0.001 infants versus adults/juveniles) but was not different between adults and juveniles (p=0.26). There was no significant effect of *sex* (F(1,40)=0.42, p=0.52) (See Supplementary table 1).

We also assessed the respiratory cycle total amplitude during the detection of Od1.1 (Fig.2 D). We observed an effect of the *phenotype* (F(1,40)=7,23, p=0.01) where blind mice display higher weight-normalized respiratory amplitude than sighted mice , no matter the age, sex or odor periods (no interactions, see supplementary table 2). In addition, we observed a global effect of *age* (F(2,80)=13.26, p<0.001), an effect of the *odor periods* (F(2,80)=9.15, p<0.001) and an interaction between *age and odor periods* (F(4,160)=42, p<0.001; Supplementary table 2). Post-hoc tests revealed that the normalized amplitude across Od1.1 *periods* did not evolve in the same way depending of *age*. At the infant age, there was a significant increase of amplitude between the pre and odor period (p<0.001) and the amplitude during the post-odor period remained high and did not return to the pre-odor level (odor vs post, p=0.28; post vs pre, p<0.001), similar to what was observed with respiratory frequency. However, this was not the case in juveniles and adults. Indeed, at the juvenile age, there was no differences between all periods (pre vs odor : p=0.11; odor vs post: p=0.99). For adults, there was no increase of amplitude between the pre and odor periods (p=0.70) but there was a significant decrease between odor and post periods (p=0.003).

Finally, there was an interaction between *age* * *sex* (F(2,80)=4.52, p=0.014) and post-hoc tests revealed no difference between male and female mice at the infant age and adult age (p=0.93 and p=0.99) but a difference between them at the juvenile age (p=0.002; Fig. 2D).

### Sniffing response in the odor habituation/cross-habituation paradigm throughout ontogeny in both blind and sighted mice

We analyzed the respiratory frequency and normalized amplitude over 20-sec time-windows following each odor presentation (20sec; Od1.1, Od1.2, Od1.3, Od1.4 and Od2) in blind and sighted mice in both males and females (Fig. 3).

**Figure 3.**
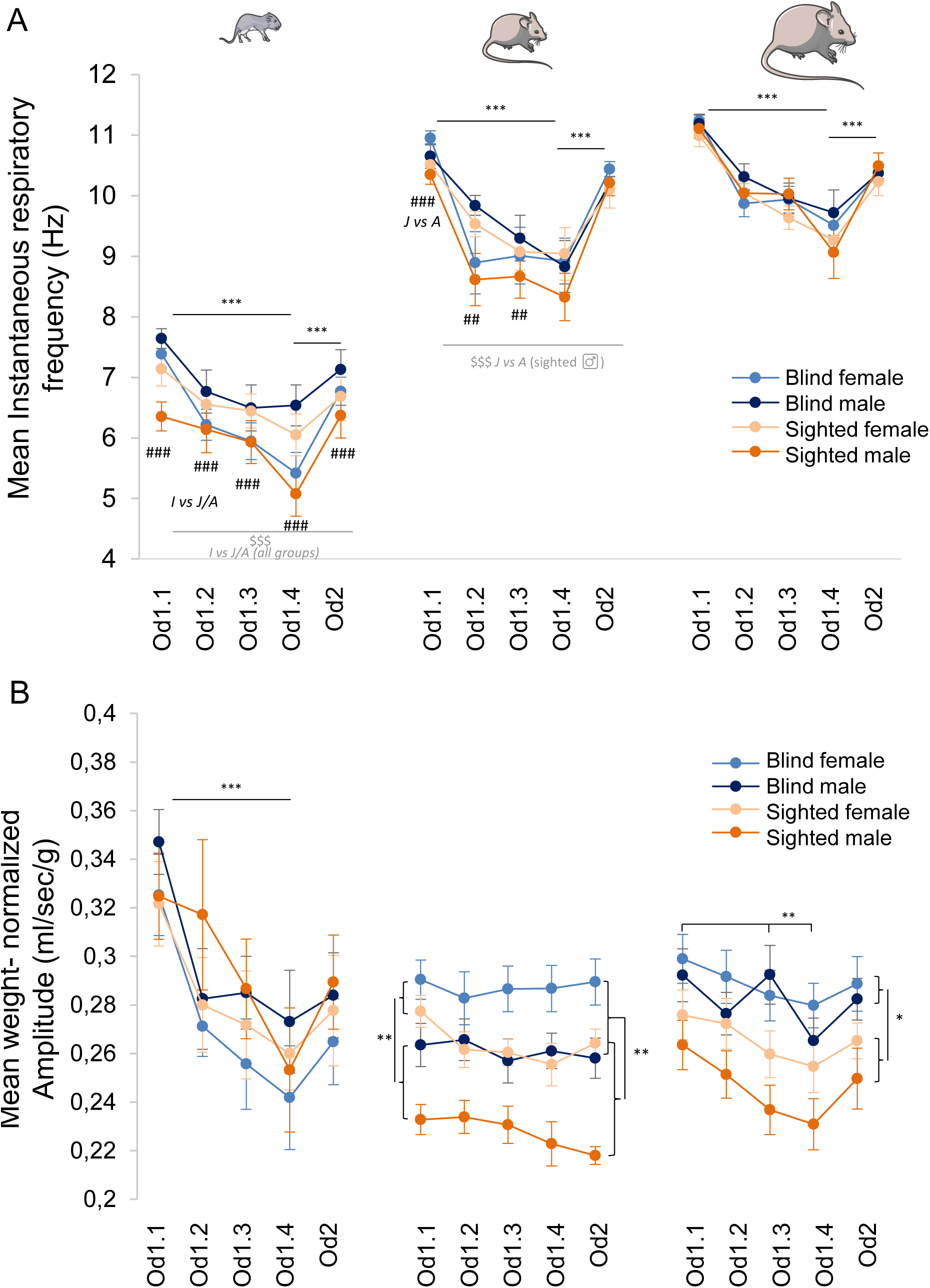
Respiratory parameters during an odor habituation/cross-habituation task throughout ontogeny in both blind and sighted mice. A- Mean instantaneous frequency of the respiratory signal (Hz) and B- Mean weight-normalized amplitude of the respiratory signal (blind female: cyan n=12; blind male: dark blue n=10; sighted female: salmon n=11; and sighted male: orange n=11). Mean +/- SEM. Repeated measures ANOVA were performed. The ANOVA was followed by Tukey post-hoc comparisons when significant. On the figure and for clarity, only difference between Od1.1 and Od1.4 and between Od1.4 and od2 are displayed as they represent an index of habituation and discrimination (*). * also displays a difference at specific age between either the sexes or phenotypes.*, p<0.05; **p<0.01, ***p<0.001. For respiratory frequency, # represents a significant difference at a specific odor presentation between ages no matter the sex or phenotype (age X odor presentations interaction; ##, p<0.01; ###, p<0.001) and $ represents a significant difference between ages for each phenotype and/or sexes (no matter the odor presentations). $$, p<0.01; $$$, p<0.001.

For the respiratory frequency (Fig. 3A), we observed a tendency for the effect of the *phenotype* (F(1,40)=3.76, p=0.06), no main effect of the *sex* (F(1,40)=0.012, p=0.91) but interactions between *phenotype* and *sex* (F(1,40)=4.59, p=0.04), between *age*, *phenotype* and *sex* (F(2,80)=3.6, p=0.032) as well as *odor presentations*, *phenotype* and *sex* (F(4,160)=2.52, p=0.043; Supplementary table 3). Further, we also observed a significant effect of *age* (F(2,80)=687.7; p<0.001) and *odor presentations* (F(4,160)=74.1 p<0.001) as well as an interaction between *age* and *odor presentations* (F(8,320)=2.5, p=0.012).

First, regarding the effect of *age* * *phenotype* * *sex*, sniffing frequency changed across ages. For all phenotype-sex combinations, except sighted males, there was a significant increase of sniffing frequency between infants and juveniles/adults but not between juveniles and adults while this was also significantly different in sighted males.

Second, regarding the effect of *odor presentations* * *age*, post-hoc analysis revealed that, for all ages, no matter the phenotype or sex, the mean respiratory frequency decreased progressively with the successive presentations of Od1 (Od1.1 vs Od1.4, p<0.001 for all ages Fig. 3A), and there was a clear cross-habituation (Od1.4 vs Od2, p<0.001 for all ages Fig. 3A). In addition, respiratory frequency increased across all ages for Od1.1; Od1.2 and Od1.3 and only between infants vs both juveniles and adults for Od1.4 and Od2.

Third, as to the *odor presentations*, *phenotype* and *sex* interaction, post-hoc tests did not reveal significant difference between phenotype/sex at each odor presentation nor in habituation/cross-habituation for each phenotype/sex when looking at the tests for Od1.1 vs other Od1 presentations and between Od1.4 and Od2.

We next analyzed the respiratory weight-normalized amplitude during the habituation/cross-habituation task (Fig. 3B; Supplementary table 4). We observed a significant effect of *phenotype* (F(1,40)=4,89, p=0.03), *age* (F(2,80)=5.96, p=0.004), *odor presentations* (F(4,160)=30.94, p<0.001) and interactions between *age and odor presentations* (F(8,320)=10.12, p<0.001), *age and sex* (F(2,80)=5.35, p=0.007), and *age and phenotype* (F(2,80)=3.43, p=0.037).

First, the *phenotype* effect was dependent on *age*, with no effect of the absence of the visual system at the infant age (p=0.99) while the difference between both phenotypes appears at the juvenile age (p=0.004) and remains significant at the adult age (p=0.03).

Second, each age has different dynamics of respiratory amplitude responses across odor presentations: at the infant age, there was a strong decrease of respiratory amplitude across repeated presentations of Od1, however there was no difference between Od1.4 and Od2 (p=0.35). At the juvenile age, the amplitude is rather stable across odor presentations (no significant differences between Od1 presentations nor between Od1.4 and Od2). For adults, albeit to a lesser degree than infants, there was also a decrease of sniffing amplitude across repeated presentations of Od1 and a tendency of increase between Od1.4 and Od2 (p=0.051).

Finally, that there was no difference between males and females at the infant and adult ages while there was one at the juvenile age (p=0.90, p= 0.0011 and p=0.72 for infants, juveniles and adults, respectively; Fig. 3B).

## Discussion

The current study provides evidence on the ontogeny of respiratory responses in a murine model of congenital blindness during an olfactory habituation/cross-habituation test. Respiration frequency has been shown to be a faithful index of olfactory performances in rats and adult mice, and we show here that it is also the case throughout ontogeny in mice. Our results also highlight the differential outcome of two respiratory parameters (i.e. frequency and amplitude) in the case of a sensory loss, in a mouse model of congenital blindness. We observed that blind mice display higher respiratory frequency and amplitude than sighted mice during detection of the first odor. For the habituation/cross-habituation, blind and sighted mice display clear habituation and discrimination using sniffing frequency as an index. The weight-normalized amplitude of the respiratory signal does not represent an index of odor habituation or discrimination throughout ontogeny. However, it highlights differences between phenotypes and sexes that were age-dependent. Notably, blind mice display higher normalized amplitude than sighted mice starting at the juvenile age regardless of the odor presentations.

### Effect of congenital blindness on odor-evoked respiratory responses throughout ontogeny

To our knowledge, this is the first study investigating the effect of congenital blindness on odor-evoked respiratory responses in developing mice. Congenital/early blindness leads to enhancement of non-visual remaining senses, such as olfactory function associated with dramatic brain reorganization (Touj et al. 2020, 2021; Zhou et al. 2017). Notably, behavioral studies pointed out that congenitally and early blind rodents have higher olfactory performances than their sighted congeners (Bouguiyoud et al. 2022, 2023; Touj et al. 2020; Zhou et al. 2017). Additionally, enhanced olfactory perception induced by post-natal visual deprivation is associated with enhancement of the power of high-frequency beta and gamma oscillations in olfactory bulb and anterior piriform cortex in rodents (Zhou et al. 2017). We therefore expected that blind mice would have a better index of olfactory performance and exhibit higher odor-evoked responses, especially in juveniles and adults as infants in both blind and sighted mice have no or only few visual experiences. Using sniffing frequency as an index of olfactory performance, we did not observe difference between blind and sighted mice in terms of olfactory abilities : odor detection, habituation, and discrimination. That might be related to the fact that our task was not challenging enough to highlight olfactory performances differences.

While olfactory performances were not impacted by the blindness, we observed that during detection of the first odor, the lack of visual inputs was associated with increased respiratory frequency and weight-normalized amplitude in mice, regardless of the age or odor period. This difference between phenotypes was observed at PN11, even if sighted infants have limited/reduced visual experience. However, it is possible that sighted PN11 mice already process some visual information through their eyelids or their eyes (eye opening for 2-day max). In addition, and interestingly, during odor habituation and discrimination, while the odor-evoked respiratory frequency was only mildly altered by congenital blindness (tendency effect only), the respiratory amplitude revealed differences between the two phenotypes starting at the juvenile age, as observed with behavioral measurements (Bouguiyoud et al., 2022). Respiratory amplitude might thus be sensitive to the accumulated visual experience.

These differences of respiratory parameters in blind mice might be related to the increased locomotor exploratory activity evidenced in several visual deprivation rodent models, even in odorless paradigm (Bouguiyoud et al. 2021; Dyer and Weldon 1975; Iura and Udo 2014; Klein and Brown 1969). Blind animals may display greater exploratory activity than sighted ones to achieve the same level of central arousal using their remaining sensory inputs (Berlyne 1955). It might also be linked to potential differences in terms of hypervigilance/attentional and motivational processes, and neural plasticity in the blind (Bouguiyoud et al. 2021, 2022; Deschênes et al. 2012; Touj et al. 2020, 2021). Thereby, when introduced in a newly environment, with or without any odor, blind mice tend to collect sensory information by increasing the sniffing frequency and the air volume intake to compensate for their blindness (Iura and Udo 2014).

In this experiment, we included both males and females without a specific underlying hypothesis on the sex effect. Regarding sniffing frequency, we did not observe a significant effect of the sex during the odor detection. There was only a significant interaction during the habituation/cross-habituation where blind males display a small-size effect difference of frequency between the juvenile and adult stage. Interestingly, for the weight-normalized amplitude, there was a significant difference between males and females at the juvenile stage. The maturation of respiratory and neuromodulatory systems as well as changes of hormonal levels might have a considerable impact on respiratory parameters (Holley et al. 2012; Patrone et al. 2020). These results shed light on the importance of considering the sex as a factor in the analysis.

### Temporal course of respiratory parameters during odor detection, habituation and discrimination throughout ontogeny in mice

In addition to the effect of blindness, we also observed major effect of the developmental age on respiratory parameters, regardless of the phenotype or sex. First regarding respiratory frequency, prior to odorant presentation, we found that infant mice displayed lower instantaneous respiratory frequency compared to juvenile or adult mice (i.e., PN10-13: 5.6Hz; PN30-33: 9.5 Hz; PN60: 9.4 Hz). Such basal respiratory parameter changes during the post-natal period have also been reported in several studies on rodents (Wang et al. 2021;Alberts and May 1980a). Globally, it seems that in mice, basal respiratory frequency increases with age, reaching a peak at the juvenile age and then stabilizing. This increase can be associated to changes in general locomotor activity across age in rodents in a novel environment (Lynn and Brown 2009; Reiber et al. 2022).

Despite this difference of respiratory frequency prior to odorant presentation, we found that, for all ages, an unfamiliar odor induces a clear increase of respiratory frequency, and this frequency is increasing with age (from 7.1 Hz to 10.6 to 11.1 for infants, juveniles and adults, respectively). Infant mice not only display lower respiratory frequency than juveniles or adults but their odor-evoked respiratory responses also have different dynamics as respiratory frequency during the post-Od1 period remained similar to the odor period. Our findings are in line with previous studies showing odor-induced sniffing behavior starting from the second week of life in developing rats (Alberts and May 1980a,b; Boulanger-Bertolus et al. 2023). Boulanger-Bertolus and colleagues (2023) recorded respiratory frequency during odor presentation in rats at PN12-15, PN22-24 and older than PN75. At the 3 ages, they showed an increase of respiratory frequency in response to the odor, and this enhancement was greater in adults compared to juveniles and infants, while no difference was observed between the two younger ages (Boulanger-Bertolus et al. 2023). Interestingly, the post-odor sniffing frequency returned to basal respiratory levels in adults and juveniles, whereas the infant sniffing response remains high. The persisting high sniffing frequency in the infants in post-odor period may be explained by the fact that infant rodents display prolonged exploratory behavior in a novel environment and towards new stimuli/objects (Bolles and Woods 1964; Campbell and Spear 1972).

In the present study, all experimental age groups displayed a habituation process toward the repeated presentation of the odor, as the respiratory frequency progressively decreased over the 4 consecutive odor presentations (Od1). To the best of our knowledge, only one study addressed the question of the ontogeny of the respiratory frequency in response to the repetitive presentation of an odor, from infant to adult rats (Boulanger-Bertolus et al. 2023). Similarly, to our data, the authors found that sniffing frequency progressively decreased across the successive trials but, in their case, faded more rapidly for younger animals.

All age groups also exhibited a clear cross-habituation, highlighted by an intense respiratory response when they were exposed to the second odor (Od2) compared to Od1.4. Those respiratory frequency responses at all ages in mice are thus comparable to what was observed in adults’ rodents (Al Koborssy et al. 2019; Coronas-Samano et al. 2016; Johnson et al. 2020; Walker et al. 2024; Wesson et al. 2008b, 2008a).

Although sniffing frequency has been used as a faithful index for odor perception (Al Koborssy et al. 2019; Boulanger-Bertolus et al. 2023; Coronas-Samano et al. 2016; Johnson et al. 2020; Walker et al. 2024), other respiratory parameters could help characterizing odor processing (Youngentob 2005; Youngentob et al. 1987). In line with the mean instantaneous respiratory frequency, infants showed an increase of respiratory amplitude after the first delivery of a new odor, which remained enhanced and did not return to the basal level in post-odor period. In juveniles and adults, the amplitude did not seem to reflect a faithful index of odor detection as the amplitude was unchanged between pre-odor and odor periods. This contrasts what was observed in adult rats (Youngentob et al. 1987). In our experiment, a ceiling effect might have occurred for juvenile and adult mice. With regards to the habituation, although a decrease was recorded in infant and adult mice across repeated presentations of Od1, it remained stable during the cross-habituation phase for both ages and appeared to be unchanged across all odor presentations in juveniles. It should be noted that the total amplitude was normalized by the weight of the animal as the weight is a confounding effect with age (Mortola et al. 1994). However, this normalization does not affect the dynamics of the response within each age. The weight-normalized respiratory amplitude is thus not an informative olfactory index throughout age, except for detection and habituation in infant mice.

## Conclusion

The analysis of odor-evoked respiratory responses, particularly during ontogeny, brings important new insight into the field of olfactory perception assessment. Changes in respiratory response is a spontaneous response that reflects olfactory performances throughout development. We demonstrated that respiratory frequency appears to be very sensitive to assess the olfactory performance independently of motor skills from the early age of development. We also showed that sniffing frequency is increasing with age and is enhanced in blind mice, compared with sighed ones. In addition, we showed that the respiratory amplitude is not a reliable index for evaluating olfactory ability across all developmental stages but can be used to discriminate both phenotypes (and sex effect) in our experimental task. Collectively, our data highlight that sniffing frequency constitutes a valid parameter to investigate both the course of odor detection, habituation, and discrimination in developing mice and that other respiratory parameters may bring complementary information of the status of the animal.

Future studies would be needed to investigate the neuronal circuits and mechanism implicated in the respiratory changes throughout development. Notably, given the importance of both respiratory frequency and amplitude on brain rhythms and synchronization (Courtiol et al. 2011; Esclassan et al. 2012; Girin et al. 2021; Juventin et al. 2023), further studies should examine whether structural plasticity may alter neuronal activity function and the role of respiration on regulation/modulation of the functional neuroplasticity, as well as the consequent behavioral changes in mouse models of congenital blindness.

## Acknowledgments

We would like to thank Ounsa Jelassi, Laurine Lacroze and Tamara Basthard-Bogain for their technical support.

The data underlying this article will be shared on reasonable request to the corresponding author.

