## Supplementary tables for "Odor-evoked respiratory responses throughout development in sighted and blind mice"

**Supplementary material : statistical tables**

**Supplementary table 1 : Sniffing frequency during the detection of Od1.1 (associated to Fig. 2C)**

| Within Subjects Effects | | | | | | | | | | | | | | | | | | | | | | | | | | | | | | | | | | | | | |
| --- | --- | --- | --- | --- | --- | --- | --- | --- | --- | --- | --- | --- | --- | --- | --- | --- | --- | --- | --- | --- | --- | --- | --- | --- | --- | --- | --- | --- | --- | --- | --- | --- | --- | --- | --- | --- | --- |
|  | | | | **Sum of Squares** | | | | | | | | | | | **df** | | | | | | | | **Mean Square** | | | | | | | | **F** | | | | | | **p** |
| **Age** | | | | | | |  | | | **1032.96810** | | |  | | | | **2** | | |  | | **516.48405** | | | | |  | | **396.24069** | | | | |  | | **3.1142e-42** |  |
| Age ✻ phenotype | | | | | | |  | | | 5.46753 | | |  | | | | 2 | | |  | | 2.73377 | | | | |  | | 2.09731 | | | | |  | | 0.12949 |  |
| Age ✻ Sex | | | | | | |  | | | 4.28937 | | |  | | | | 2 | | |  | | 2.14469 | | | | |  | | 1.64538 | | | | |  | | 0.19940 |  |
| Age ✻ phenotype ✻ Sex | | | | | | |  | | | 0.34924 | | |  | | | | 2 | | |  | | 0.17462 | | | | |  | | 0.13397 | | | | |  | | 0.87482 |  |
| Residual | | | | | | |  | | | 104.27683 | | |  | | | | 80 | | |  | | 1.30346 | | | | |  | |  | | | | |  | |  |  |
| **od1.1 period** | | | | | | |  | | | **137.41847** | | |  | | | | **2** | | |  | | **68.70924** | | | | |  | | **88.49427** | | | | |  | | **5.3340e-21** |  |
| od1.1 period ✻ phenotype | | | | | | |  | | | 2.98337 | | |  | | | | 2 | | |  | | 1.49168 | | | | |  | | 1.92122 | | | | |  | | 0.15312 |  |
| od1.1 period ✻ Sex | | | | | | |  | | | 2.94552 | | |  | | | | 2 | | |  | | 1.47276 | | | | |  | | 1.89685 | | | | |  | | 0.15673 |  |
| od1.1 period ✻ phenotype ✻ Sex | | | | | | |  | | | 0.20947 | | |  | | | | 2 | | |  | | 0.10473 | | | | |  | | 0.13489 | | | | |  | | 0.87401 |  |
| Residual | | | | | | |  | | | 62.11407 | | |  | | | | 80 | | |  | | 0.77643 | | | | |  | |  | | | | |  | |  |  |
| **Age ✻ od1.1 period** | | | | | | |  | | | **30.24529** | | |  | | | | **4** | | |  | | **7.56132** | | | | |  | | **10.48760** | | | | |  | | **1.4321e0-7** |  |
| Age ✻ od1.1 period ✻ phenotype | | | | | | |  | | | 1.28921 | | |  | | | | 4 | | |  | | 0.32230 | | | | |  | | 0.44704 | | | | |  | | 0.77444 |  |
| Age ✻ od1.1 period ✻ Sex | | | | | | |  | | | 3.90587 | | |  | | | | 4 | | |  | | 0.97647 | | | | |  | | 1.35436 | | | | |  | | 0.25223 |  |
| Age ✻ od1.1 period ✻ phenotype ✻ Sex | | | | | | |  | | | 4.09770 | | |  | | | | 4 | | |  | | 1.02442 | | | | |  | | 1.42088 | | | | |  | | 0.22941 |  |
| Residual | | | | | | |  | | | 115.35641 | | |  | | | | 160 | | |  | | 0.72098 | | | | |  | |  | | | | |  | |  |  |
| Between Subjects Effects | | | | | | | | | | | | | | | | | | | | | | | | | | | | | | | | |  |  |  |  |  |
|  | | **Sum of Squares** | | | | **df** | | | | | **Mean Square** | | | | | | | **F** | | | | | | | **p** | | | | | | | |  |  |  |  |  |
| **phenotype** |  | 19.73738 |  | | | 1 | |  | | | 19.73738 | | |  | | | | 10.37808 | | | | | |  | **0.0025357** | | | | |  | | |  |  |  |  |  |
| Sex |  | 0.80117 |  | | | 1 | |  | | | 0.80117 | | |  | | | | 0.42126 | | | | | |  | 0.5200173 | | | | |  | | |  |  |  |  |  |
| phenotype ✻ Sex |  | 2.85981 |  | | | 1 | |  | | | 2.85981 | | |  | | | | 1.50371 | | | | | |  | 0.2272711 | | | | |  | | |  |  |  |  |  |
| Residual |  | 76.07333 |  | | | 40 | |  | | | 1.90183 | | |  | | | |  | | | | | |  |  | | | | |  | | |  |  |  |  |  |

**Within-subject factors:**

- Age (infants / juveniles / adults)
- Od1.1 period (pre / odor / post)

**Between-subject factors:**

- Phenotype (blind / sighted)
- Sex (female / male)

**Supplementary table 2 : Normalized respiratory amplitude during detection of odor1.1 (associated to Fig. 2D)**

| Within Subjects Effects | | | | | | | | | | | | | | | | | | | | | | |
| --- | --- | --- | --- | --- | --- | --- | --- | --- | --- | --- | --- | --- | --- | --- | --- | --- | --- | --- | --- | --- | --- | --- |
|  | | | | | **Sum of Squares** | | | | | **df** | | | | **Mean Square** | | | | **F** | | | | **p** |
| **Age** | | |  | | **0.0947098** | | | |  | **2** | |  | | **0.0473549** | | |  | **13.261369** | | |  | **1.0615e0-5** |
| Age ✻ phenotype | | |  | | 0.0010115 | | | |  | 2 | |  | | 5.0575e-4 | | |  | 0.141633 | | |  | 0.868157 |
| **Age ✻ Sex** | | |  | | **0.0322623** | | | |  | **2** | |  | | **0.0161312** | | |  | **4.517404** | | |  | **0.013842** |
| Age ✻ phenotype ✻ Sex | | |  | | 6.4998e-4 | | | |  | 2 | |  | | 3.2499e-4 | | |  | 0.091011 | | |  | 0.913102 |
| Residual | | |  | | 0.2856712 | | | |  | 80 | |  | | 0.0035709 | | |  |  | | |  |  |
| **od1.1 period** | | |  | | **0.0117904** | | | |  | **2** | |  | | **0.0058952** | | |  | **9.149052** | | |  | **2.6412e0-4** |
| od1.1 period ✻ phenotype | | |  | | 7.1218e-4 | | | |  | 2 | |  | | 3.5609e-4 | | |  | 0.552636 | | |  | 0.577612 |
| od1.1 period ✻ Sex | | |  | | 0.0017227 | | | |  | 2 | |  | | 8.6136e-4 | | |  | 1.336789 | | |  | 0.268491 |
| od1.1 period ✻ phenotype ✻ Sex | | |  | | 2.7162e-4 | | | |  | 2 | |  | | 1.3581e-4 | | |  | 0.210769 | | |  | 0.810409 |
| Residual | | |  | | 0.0515481 | | | |  | 80 | |  | | 6.4435e-4 | | |  |  | | |  |  |
| **Age ✻ od1.1 period** | | |  | | **0.0684489** | | | |  | **4** | |  | | **0.0171122** | | |  | **42.002126** | | |  | **4.8062e-24** |
| Age ✻ od1.1 period ✻ phenotype | | |  | | 0.0015426 | | | |  | 4 | |  | | 3.8566e-4 | | |  | 0.946597 | | |  | 0.438664 |
| Age ✻ od1.1 period ✻ Sex | | |  | | 0.0038998 | | | |  | 4 | |  | | 9.7494e-4 | | |  | 2.392996 | | |  | 0.052830 |
| Age ✻ od1.1 period ✻ phenotype ✻ Sex | | |  | | 0.0011377 | | | |  | 4 | |  | | 2.8442e-4 | | |  | 0.698118 | | |  | 0.594320 |
| Residual | | |  | | 0.0651861 | | | |  | 160 | |  | | 4.0741e-4 | | |  |  | | |  |  |
| Between Subjects Effects | | | | | | | | | | | | | | | | | | | |  |  |  |
|  | | **Sum of Squares** | | | | **df** | | **Mean Square** | | | | | **F** | | | **p** | | | |  |  |  |
| **phenotype** |  | **0.0400180** | |  | | **1** |  | **0.0400180** | | |  | | **7.22937** | |  | **0.010406** | | |  |  |  |  |
| Sex |  | 0.0054192 | |  | | 1 |  | 0.0054192 | | |  | | 0.97900 | |  | 0.328394 | | |  |  |  |  |
| phenotype ✻ Sex |  | 0.0033344 | |  | | 1 |  | 0.0033344 | | |  | | 0.60236 | |  | 0.442244 | | |  |  |  |  |
| Residual |  | 0.2214187 | |  | | 40 |  | 0.0055355 | | |  | |  | |  |  | | |  |  |  |  |

**Within-subject factors:**

- Age (infants / juveniles / adults)
- Od1.1 period (pre / odor / post)

**Between-subject factors:**

- Phenotype (blind / sighted)
- Sex (female / male)

**Supplementary table 3 : Sniffing frequency during habituation/cross-habituation (associated to Fig. 3A)**

| Within Subjects Effects | | | | | | | | | | | | | | | | | | | | | | | |
| --- | --- | --- | --- | --- | --- | --- | --- | --- | --- | --- | --- | --- | --- | --- | --- | --- | --- | --- | --- | --- | --- | --- | --- |
|  | | | | | | **Sum of Squares** | | | | | **df** | | | | **Mean Square** | | | | **F** | | | | **p** |
| **age** | | | | |  | **1746.78262** | | | |  | **2** | | |  | **873.39131** | | |  | **687.76265** | | |  | **4.0069e-51** |
| age ✻ phenotype | | | | |  | 1.07599 | | | |  | 2 | | |  | 0.53799 | | |  | 0.42365 | | |  | 0.656113 |
| age ✻ sex | | | | |  | 1.93060 | | | |  | 2 | | |  | 0.96530 | | |  | 0.76014 | | |  | 0.470950 |
| **age ✻ phenotype ✻ sex** | | | | |  | **9.15451** | | | |  | **2** | | |  | **4.57726** | | |  | **3.60442** | | |  | **0.031709** |
| Residual | | | | |  | 101.59218 | | | |  | 80 | | |  | 1.26990 | | |  |  | | |  |  |
| **odor presentations** | | | | |  | **219.64574** | | | |  | **4** | | |  | **54.91144** | | |  | **74.06965** | | |  | **2.0655e-35** |
| odor presentations ✻ phenotype | | | | |  | 2.12540 | | | |  | 4 | | |  | 0.53135 | | |  | 0.71674 | | |  | 0.581651 |
| odor presentations ✻ sex | | | | |  | 1.60751 | | | |  | 4 | | |  | 0.40188 | | |  | 0.54209 | | |  | 0.705039 |
| **odor presentations ✻ phenotype ✻ sex** | | | | |  | **7.47298** | | | |  | **4** | | |  | **1.86825** | | |  | **2.52006** | | |  | **0.043282** |
| Residual | | | | |  | 118.61579 | | | |  | 160 | | |  | 0.74135 | | |  |  | | |  |  |
| **age ✻ odor presentations** | | | | |  | **14.05335** | | | |  | **8** | | |  | **1.75667** | | |  | **2.50265** | | |  | **0.011958** |
| age ✻ odor presentations ✻ phenotype | | | | |  | 3.04297 | | | |  | 8 | | |  | 0.38037 | | |  | 0.54190 | | |  | 0.824605 |
| age ✻ odor presentations ✻ sex | | | | |  | 0.94372 | | | |  | 8 | | |  | 0.11796 | | |  | 0.16806 | | |  | 0.994872 |
| age ✻ odor presentations ✻ phenotype ✻ sex | | | | |  | 5.64167 | | | |  | 8 | | |  | 0.70521 | | |  | 1.00468 | | |  | 0.432275 |
| Residual | | | | |  | 224.61551 | | | |  | 320 | | |  | 0.70192 | | |  |  | | |  |  |
| Between Subjects Effects | | | | | | | | | | | | | | | | | | | | |  |  |  |
|  | | | **Sum of Squares** | | | | **df** | | **Mean Square** | | | | **F** | | | | **p** | | | |  |  |  |
| phenotype | |  | 10.981863 |  | | | 1 |  | 10.981863 | | |  | 3.756888 | | |  | 0.059671 | | |  |  |  |  |
| sex |  | | 0.034531 |  | | | 1 |  | 0.034531 | | |  | 0.011813 | | |  | 0.913994 | | |  |  |  |  |
| **phenotype ✻ sex** |  | | **13.416351** |  | | | **1** |  | **13.416351** | | |  | **4.589725** | | |  | **0.038304** | | |  |  |  |  |
| Residual |  | | 116.925110 |  | | | 40 |  | 2.923128 | | |  |  | | |  |  | | |  |  |  |  |

**Within-subject factors:**

- Age (infants / juveniles / adults)
- Odor presentations (odor period of Od1.1, Od1.2, Od1.3, Od1.4 & Od2)

**Between-subject factors:**

- Phenotype (blind / sighted)
- Sex (female / male)

**Supplementary table 4 : Normalized respiratory amplitude during habituation/cross-habituation (associated to Fig. 3B)**

| Within Subjects Effects | | | | | | | | | | | | | | | | | | | | | | |
| --- | --- | --- | --- | --- | --- | --- | --- | --- | --- | --- | --- | --- | --- | --- | --- | --- | --- | --- | --- | --- | --- | --- |
|  | | | | **Sum of Squares** | | | | | | **df** | | | **Mean Square** | | | | | **F** | | | | **p** |
| **age** | | |  | **0.0730553** | | | | |  | **2** |  | | **0.0365277** | | | |  | **5.9575732** | | |  | **0.0038736** |
| **age ✻ phenotype** | | |  | **0.0420811** | | | | |  | **2** |  | | **0.0210405** | | | |  | **3.4316618** | | |  | **0.0371657** |
| **age ✻ sex** | | |  | **0.0655951** | | | | |  | **2** |  | | **0.0327975** | | | |  | **5.3491988** | | |  | **0.0066011** |
| age ✻ phenotype ✻ sex | | |  | 3.7728e-5 | | | | |  | 2 |  | | 1.8864e-5 | | | |  | 0.0030767 | | |  | 0.9969282 |
| Residual | | |  | 0.4905039 | | | | |  | 80 |  | | 0.0061313 | | | |  |  | | |  |  |
| **odor presentations** | | |  | **0.0904434** | | | | |  | **4** |  | | **0.0226108** | | | |  | **30.9404665** | | |  | **4.4491e-19** |
| odor presentations ✻ phenotype | | |  | 0.0033180 | | | | |  | 4 |  | | 8.2950e-4 | | | |  | 1.1350752 | | |  | 0.3419664 |
| odor presentations ✻ sex | | |  | 0.0015807 | | | | |  | 4 |  | | 3.9517e-4 | | | |  | 0.5407451 | | |  | 0.7060173 |
| odor presentations ✻ phenotype ✻ sex | | |  | 0.0023081 | | | | |  | 4 |  | | 5.7703e-4 | | | |  | 0.7895964 | | |  | 0.5335219 |
| Residual | | |  | 0.1169257 | | | | |  | 160 |  | | 7.3079e-4 | | | |  |  | | |  |  |
| **age ✻ odor presentations** | | |  | **0.0544617** | | | | |  | **8** |  | | **0.0068077** | | | |  | **10.1205770** | | |  | **1.3345e-12** |
| age ✻ odor presentations ✻ phenotype | | |  | 0.0071250 | | | | |  | 8 |  | | 8.9062e-4 | | | |  | 1.3240248 | | |  | 0.2305427 |
| age ✻ odor presentations ✻ sex | | |  | 0.0027775 | | | | |  | 8 |  | | 3.4719e-4 | | | |  | 0.5161403 | | |  | 0.8441800 |
| age ✻ odor presentations ✻ phenotype ✻ sex | | |  | 0.0054616 | | | | |  | 8 |  | | 6.8270e-4 | | | |  | 1.0149216 | | |  | 0.4244489 |
| Residual | | |  | 0.2152515 | | | | |  | 320 |  | | 6.7266e-4 | | | |  |  | | |  |  |
| Between Subjects Effects | | | | | | | | | | | | | | | | | | | |  |  |  |
|  | | **Sum of Squares** | | | | **df** | | **Mean Square** | | | | | | **F** | | **p** | | | |  |  |  |
| **phenotype** |  | **0.0501279** | | |  | **1** |  | **0.0501279** | | | |  | | **4.88660** |  | **0.032848** | | |  |  |  |  |
| sex |  | 0.0131187 | | |  | 1 |  | 0.0131187 | | | |  | | 1.27884 |  | 0.264849 | | |  |  |  |  |
| phenotype ✻ sex |  | 0.0048853 | | |  | 1 |  | 0.0048853 | | | |  | | 0.47624 |  | 0.494117 | | |  |  |  |  |
| Residual |  | 0.4103295 | | |  | 40 |  | 0.0102582 | | | |  | |  |  |  | | |  |  |  |  |

**Within-subject factors:**

- Age (infants / juveniles / adults)
- Odor presentations (odor period of Od1.1, Od1.2, Od1.3, Od1.4 & Od2)

**Between-subject factors:**

- Phenotype (blind / sighted)
- Sex (female / male)
